## Supplementary Data for "Multi-scale dynamic imaging reveals that cooperative motility behaviors promote efficient predation in bacteria"

### Table of contents

Supplementary Figure 1 **2**

Supplementary Figure 2 **3**

Supplementary Figure 3 **4**

Supplementary Figure 4 **5**

Supplementary Figure 5 **7**

**Supplementary Table S1. 8**

#### Supplementary Figure 1

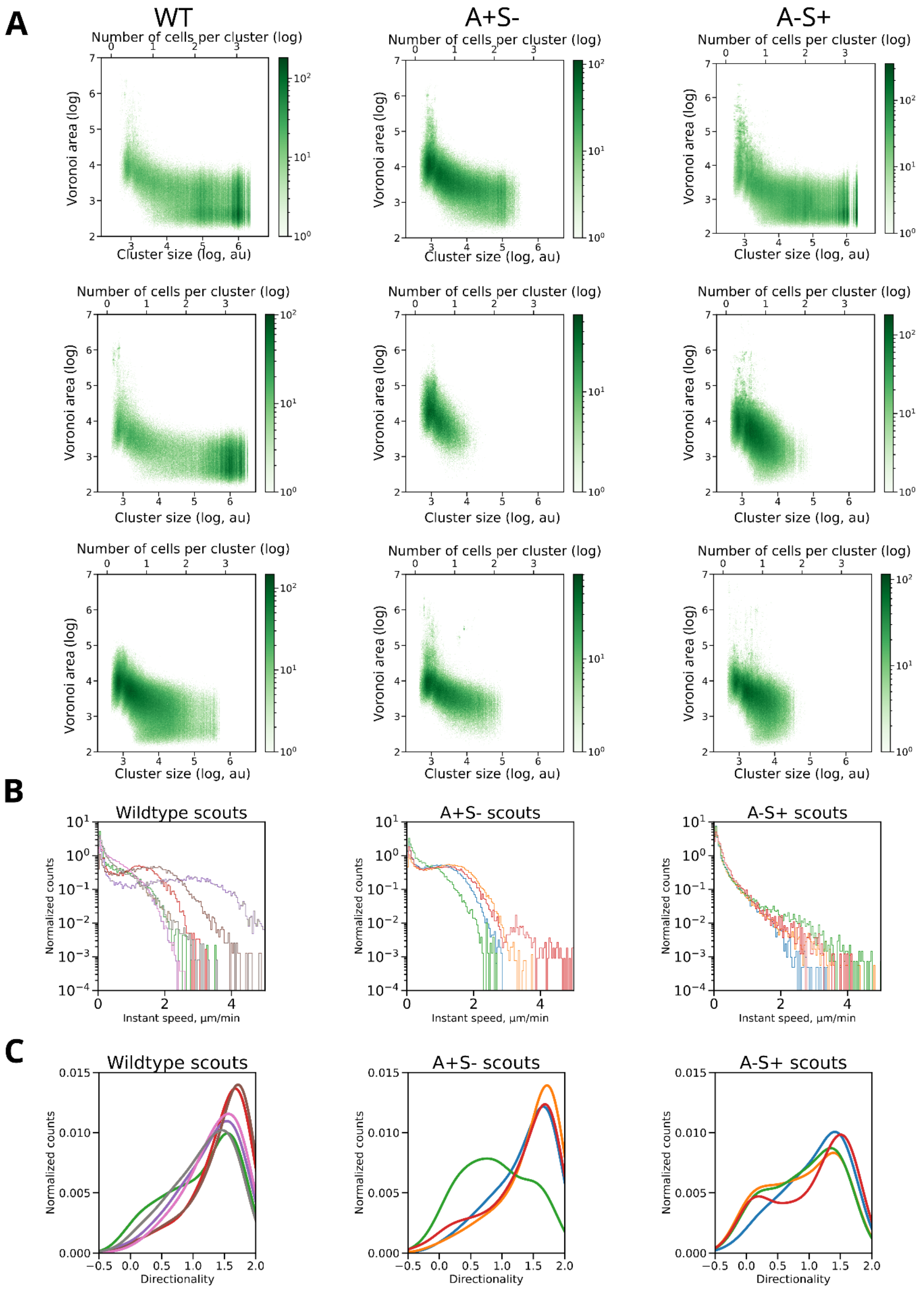


A) Examples of 2D Voronoi area-cluster size histograms for wild-type, S-motile cells (A-S+) and A-motile (A+S-) cells from experimental replicates (left, middle and right columns, respectively).

B) Histograms of instantaneous speed of scout cells from experimental replicates used in the main figure for wild-type, S-motile cells (A-S+) and A-motile (A+S-) cells. While isolated and small groups of S-motile cells can be found ahead of the predation front, A-motility is required for their mobility in regions void of prey.

C) Histograms of directionality of scout cells from experimental replicates used in the main figure for wild-type, S-motile cells (A-S+) and A-motile (A+S-) cells.

**Supplementary Figure 2**

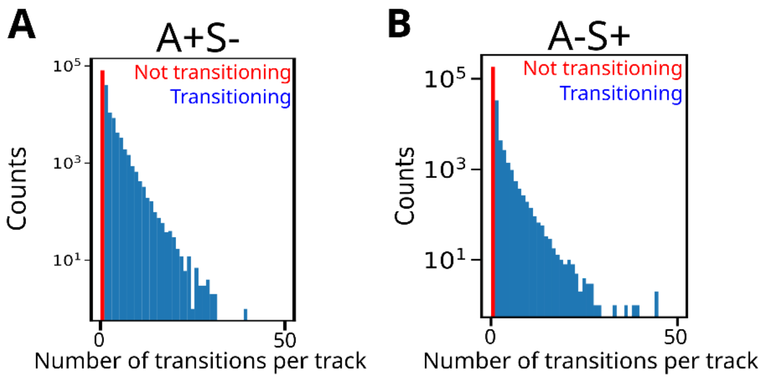


A) Histogram of number of transitions per trajectory in A-motile cells (A+S-).

B) Histogram of number of transitions per trajectory in S-motile cells (A-S+).

**Supplementary Figure 3**

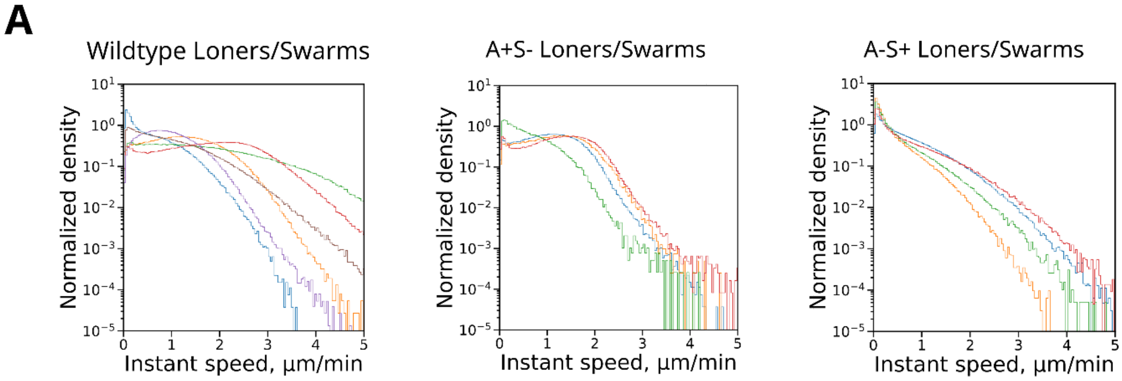


A) Histograms of instantaneous speed of swarm cells from experimental replicates used in the main figure for wild-type, S-motile cells (A-S+) and A-motile (A+S-) cells.

**Supplementary Figure 4**

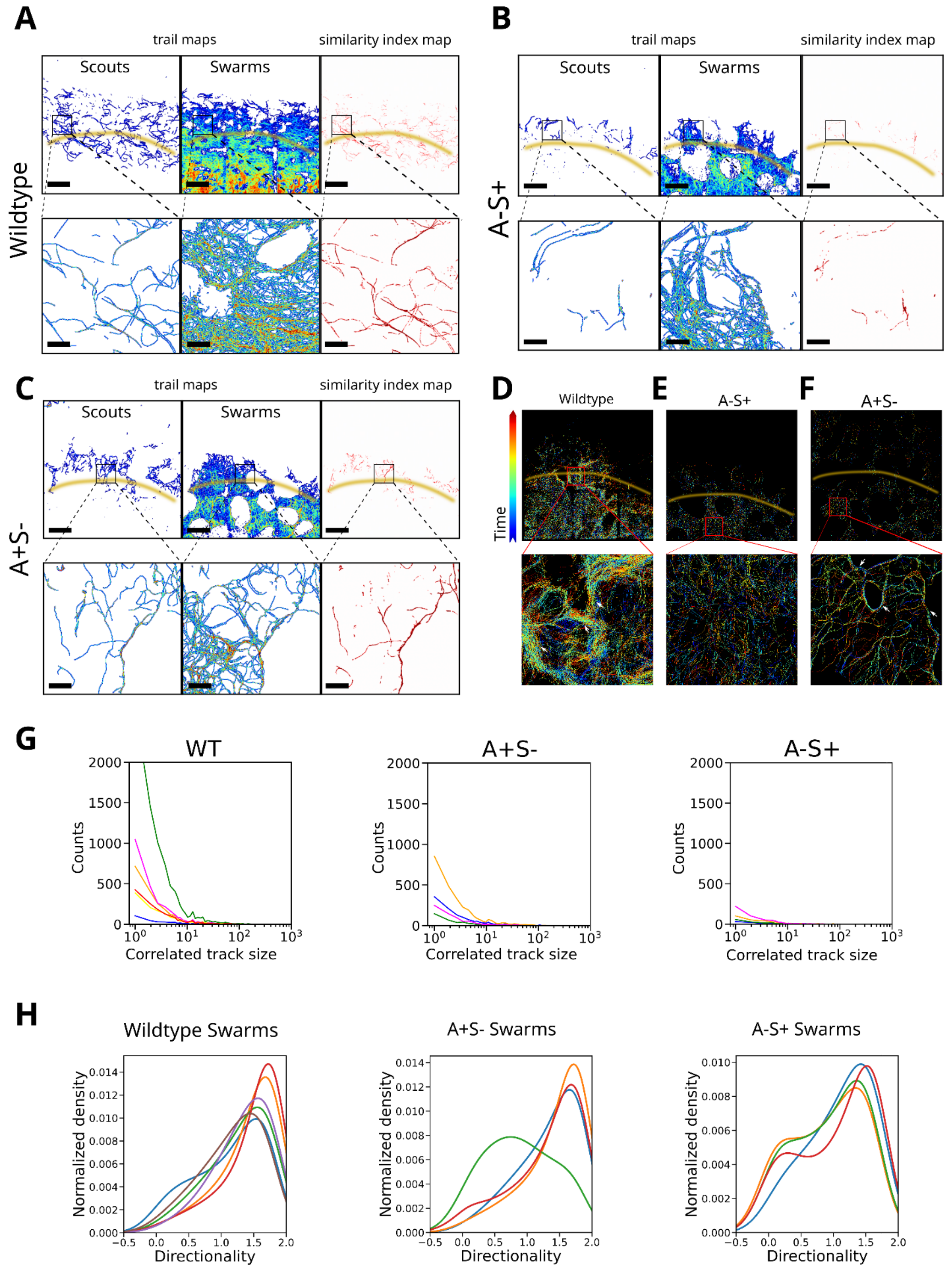


A-C) Examples of trail maps of scout and swarm cells and similarity index map for wildtype (A), A-S+ (B) and A+S- cells (C). Yellow lines delimitate the predation front. Scalebars = 100 µm. Scalebars of zoomed boxed areas = 20 µm.

D-F) Examples of overlays of all trajectories for wildtype (D), S-motile (A-S+) cells (E) and A-motile (A+S-) cells (F). Yellow lines delimitate the predation front. White arrows in the zoom of the boxed areas point to examples of trails.

G) Histogram of length of overlapping tracks (Similarity track size) used in the main figure for wild-type, S-motile cells (A-S+) and A-motile (A+S-) cells.

H) Histogram of swarm cells tracks directionality used in the main figure for wild-type, S-motile cells (A-S+) and A-motile (A+S-) cells.

**Supplementary Figure 5**

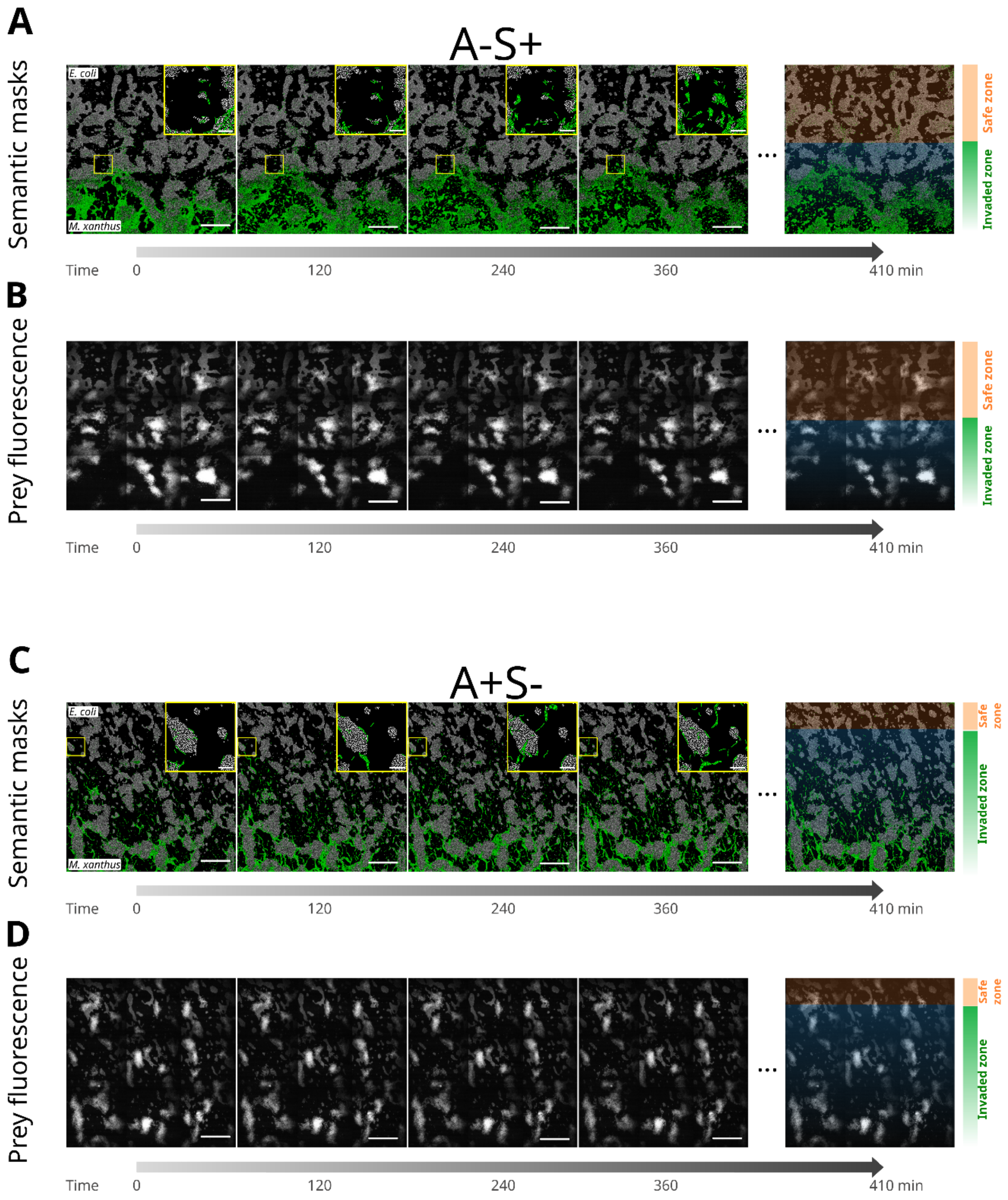


A) Evolution of predation over time at the predation forefront of S-motile cells (A-S+) visualized with semantically segmented large ROI containing the masks for *M. xanthus* (green) and *E. coli* (white). Two zones were defined: i) where no invasion occurred (orange shaded area) and ii) where *M. xanthus* cells invaded *E. coli* (blue shaded area). Scalebars = 100 µm. Scalebars of zoomed images = 20 µm

B) Evolution of the total raw fluorescence signal from *E. coli* cells over time invaded by S-motile *M. xanthus* cells (A-S+). Orange and blue boxed areas highlight the safe zone, where no predation occurred and the predation zone, where active predation is occurring, respectively. Scalebars = 100 µm. Scalebars of zoomed images = 20 µm

C-D) same as panels A and B but for A-motile *M. xanthus* cells (A+S-)

**Supplementary Table S1**

Bacterial strains used in this study.

| **Strain** | **Genotype** | **Strain origin** |
|---|---|---|
| *E. coli* | MG1655 wildtype | |
| *E. coli* HU-mCherry | MG1655 HU-mCherry | Espeli laboratory collection |
| *M. xanthus* | DZ2 wildtype | Mignot laboratory collection |
| *M. xanthus* cytosolic-sfGFP | DZ2 pSWU19-Pm1000-sfGFP | Mignot laboratory collection |
| *M. xanthus* A+S- OMss-sfGFP | DZ2 ΩpilA pSWU19-PpilA-OMss-sfGFP | Mignot laboratory collection |
| *M. xanthus* A-S+ OMss-sfGFP | DZ2 GltJ DNterm222 pSWU19-PpilA-OMss-sfGFP | Mignot laboratory collection |
| *M. xanthus* A+S- OMss-mCherry | DZ2 DpilA pSWU19-PpilA-OMss-mCherry | Mignot laboratory collection |
| *M. xanthus* AglZ-NeonGreen | Allelic replacement of *aglZ* by *aglZ-NeonGreen* | Mignot laboratory collection |
